## Supplementary Information for "Hip stabilization in an australopithecine-like hip: the influence of shape on muscle activation"

### Supplemental Information

#### Kinematic comparison to previous work

Joint angles are shown in the following five figures.  All trials for the ADL human-like configuration are shown with blue dashed lines. The average is shown with a solid blue line while the values from Winter [74] at natural cadence are shown with a solid black line (where applicable).

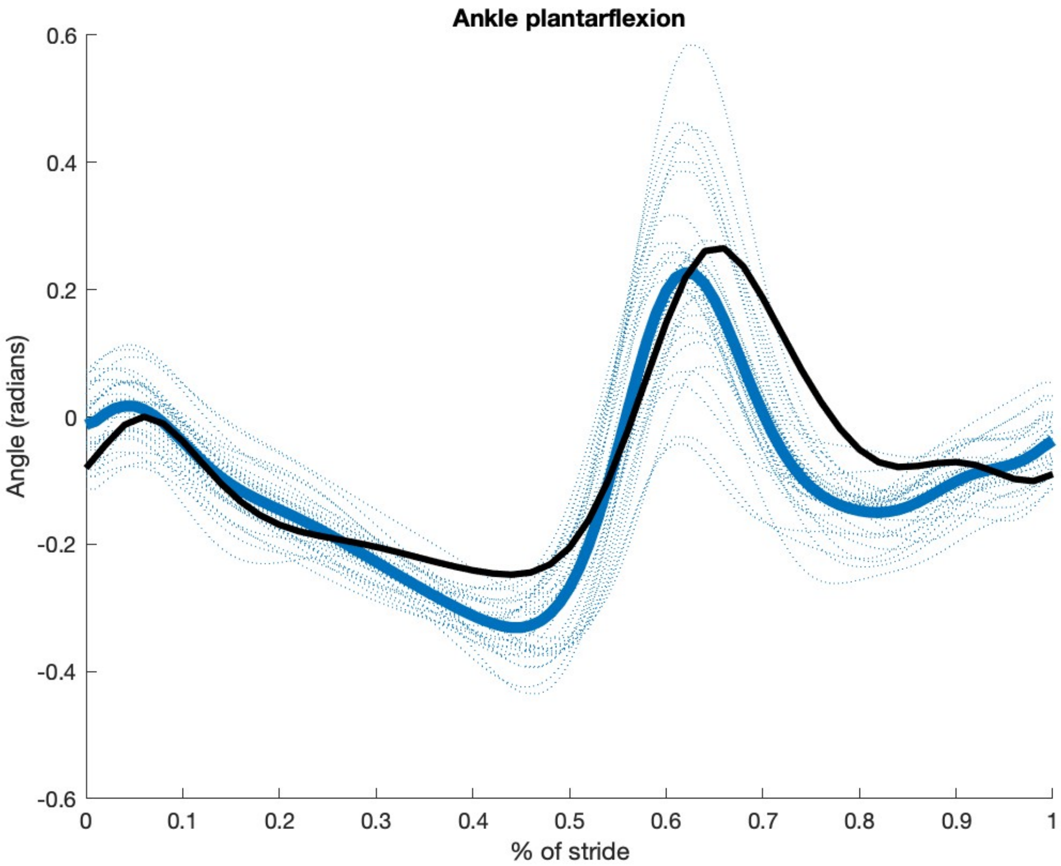

Ankle plantarflexion
Angle (radians)
0.6
0.4
0.2
0
-0.2
-0.4
-0.6
0
0.1
0.2
0.3
0.4
0.5
0.6
0.7
0.8
0.9
1
% of stride

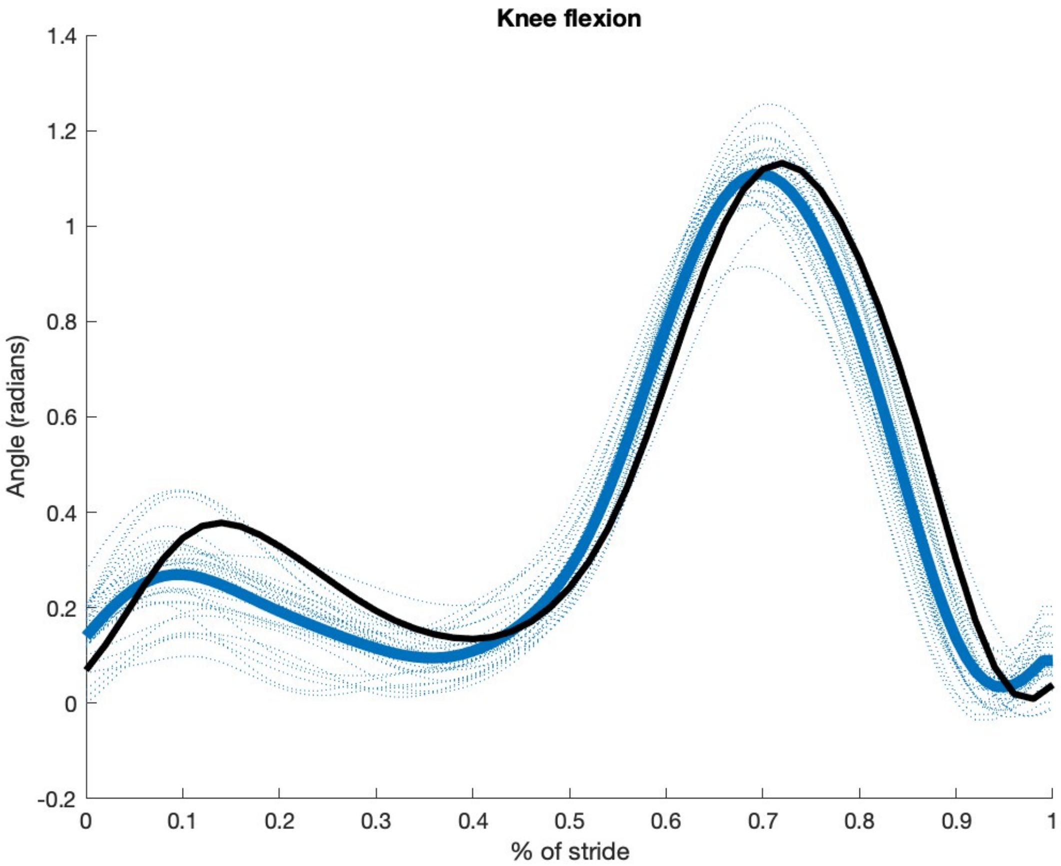

Knee flexion
Angle (radians)
% of stride
1.4
1.2
1
0.8
0.6
0.4
0.2
0
-0.2
0
0.1
0.2
0.3
0.4
0.5
0.6
0.7
0.8
0.9
1

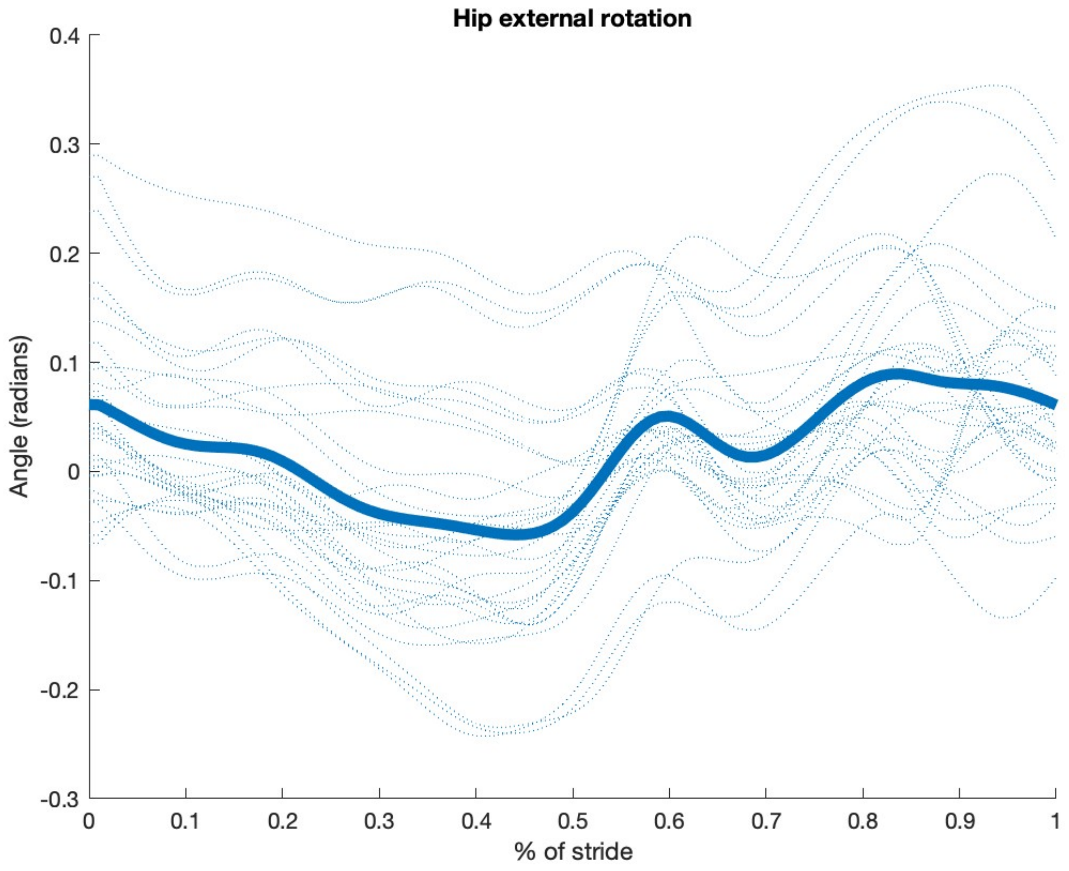

Hip external rotation
Angle (radians)
% of stride
0.4
0.3
0.2
0.1
0
-0.1
-0.2
-0.3
0
0.1
0.2
0.3
0.4
0.5
0.6
0.7
0.8
0.9
1

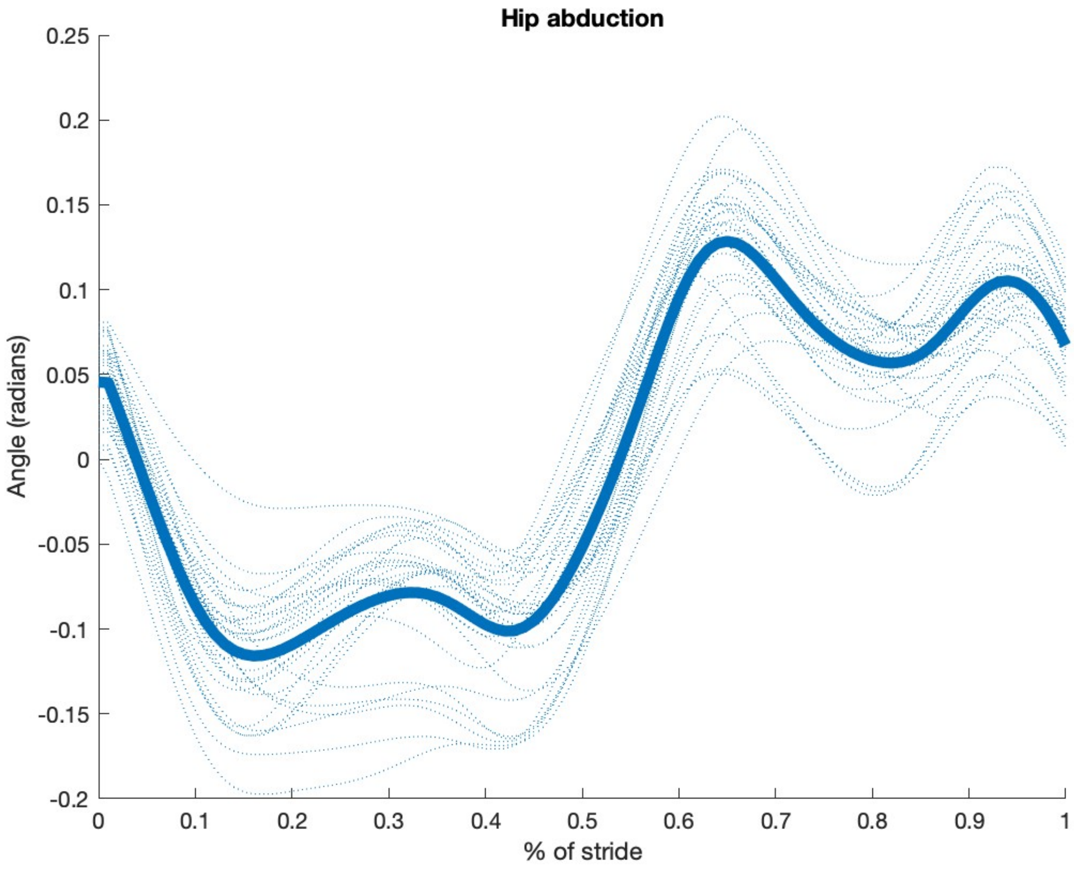

Hip abduction
Angle (radians)
0.25
0.2
0.15
0.1
0.05
0
-0.05
-0.1
-0.15
-0.2
0
0.1
0.2
0.3
0.4
0.5
0.6
0.7
0.8
0.9
1
% of stride

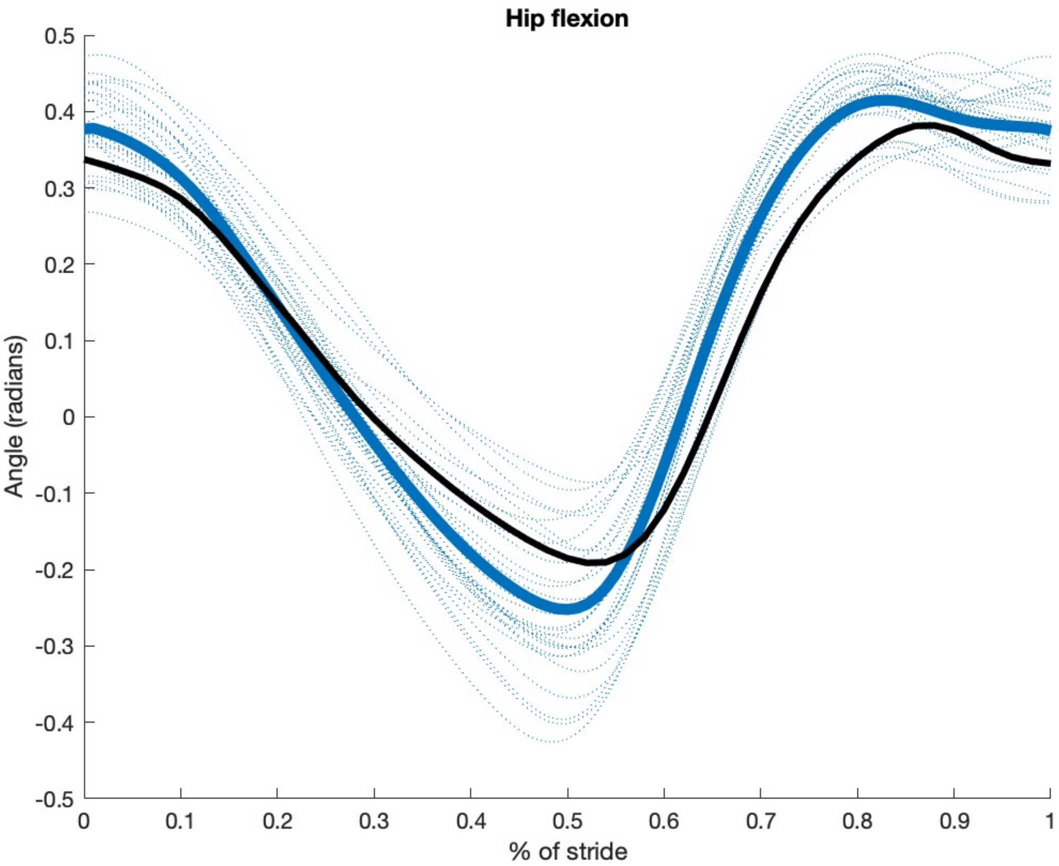

Hip flexion
Angle (radians)
% of stride
0.5
0.4
0.3
0.2
0.1
0
-0.1
-0.2
-0.3
-0.4
-0.5
0
0.1
0.2
0.3
0.4
0.5
0.6
0.7
0.8
0.9
1

**Gluteal muscle activation compared to previous work**

Activation of the gluteal muscles compared to previously published data [64]. All trials for the ADL human-like configuration are shown with blue dashed lines. The average is shown with a solid blue line while the values from Sylvester et al. are shown with a solid black line.

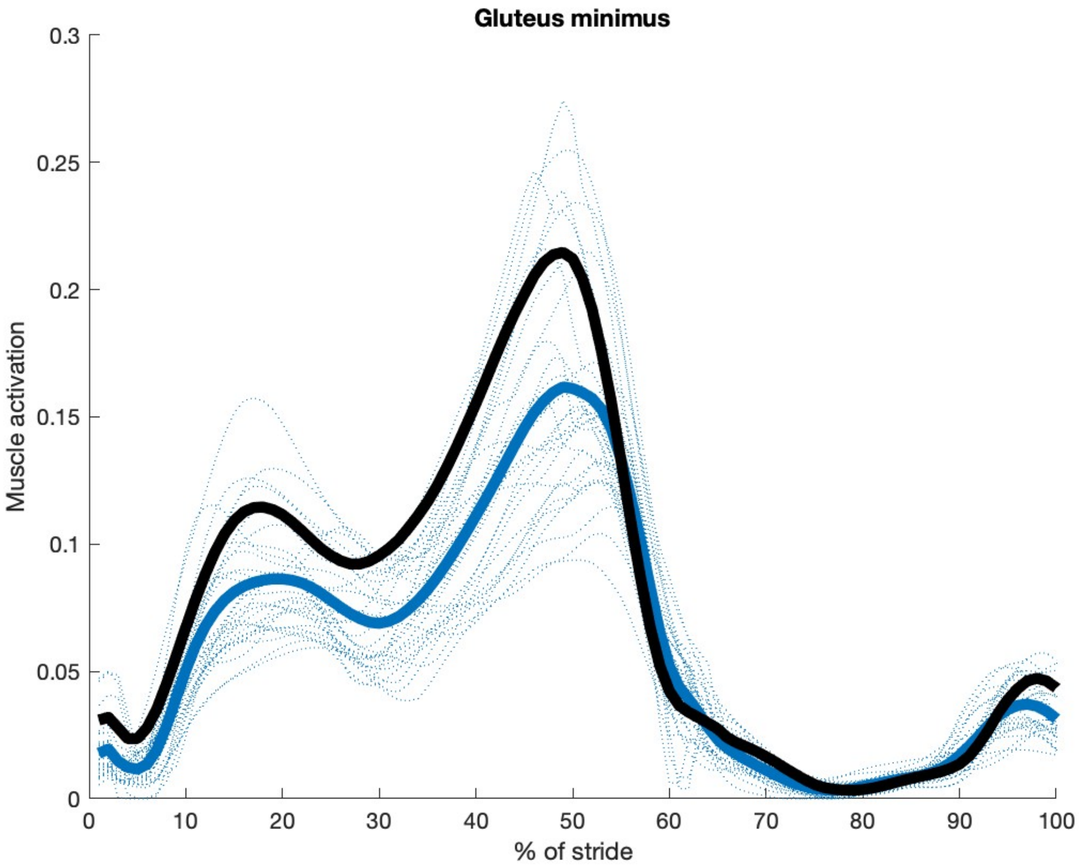

Gluteus minimus
Muscle activation
0.3
0.25
0.2
0.15
0.1
0.05
0
0
10
20
30
40
50
60
70
80
90
100
% of stride

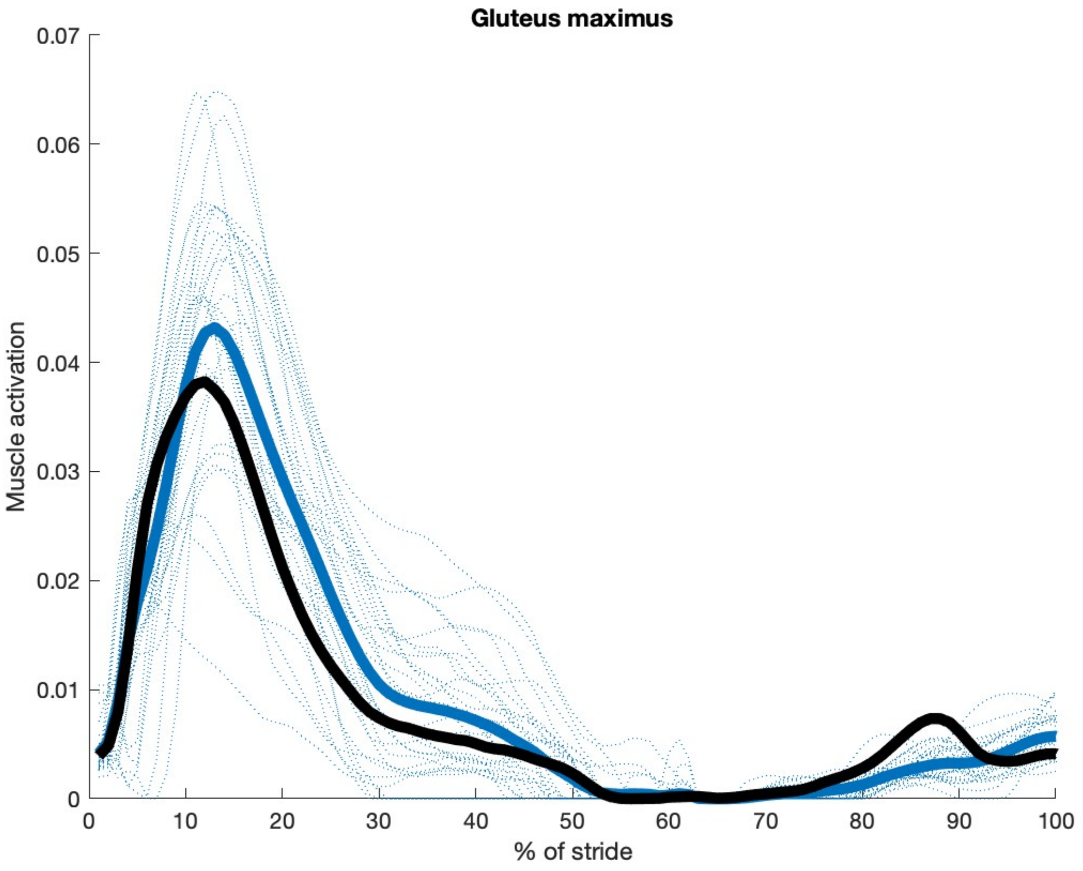

Gluteus maximus
Muscle activation
0.07
0.06
0.05
0.04
0.03
0.02
0.01
0
0
10
20
30
40
50
60
70
80
90
100
% of stride

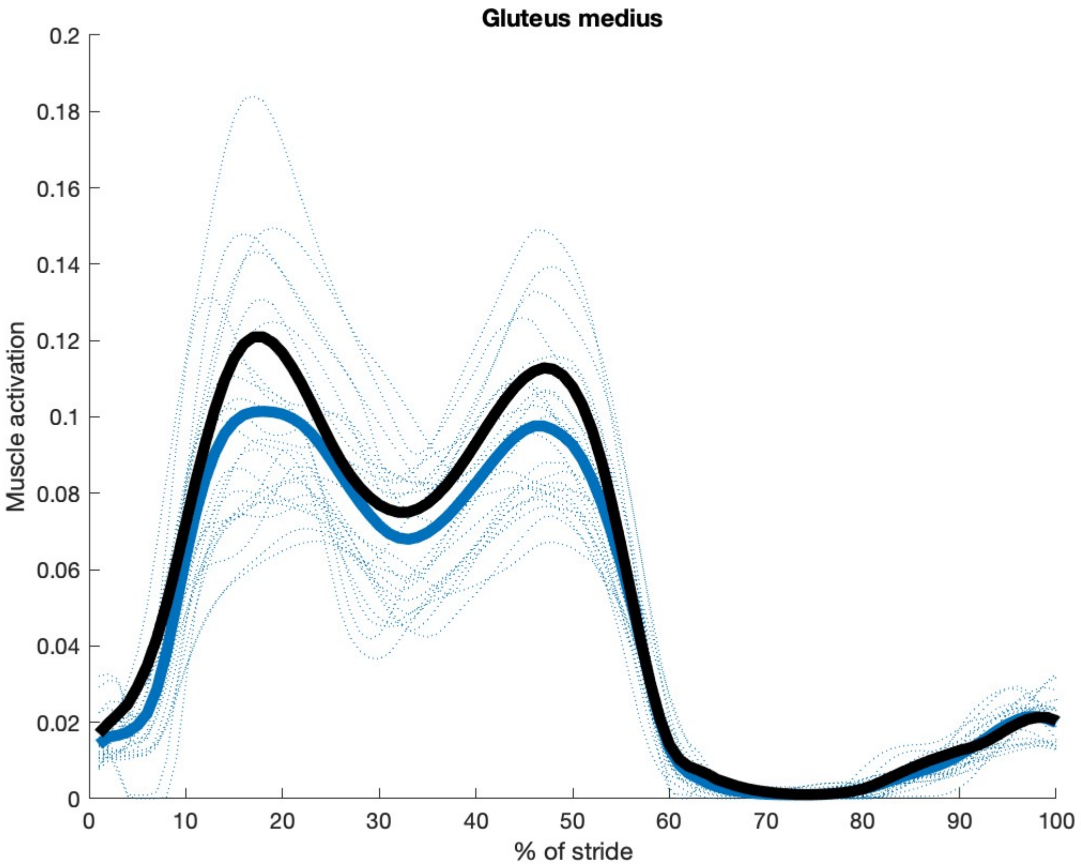
Gluteus medius
Muscle activation
0.2
0.18
0.16
0.14
0.12
0.1
0.08
0.06
0.04
0.02
0
0
10
20
30
40
50
60
70
80
90
100
% of stride

**Muscle element moment arms**

The moment arms over which forces act about joints can be useful in visualizing and evaluating the relative effects of various parts of a structure (such as the hip) in static analyses. Consequently, important early research used them in comparisons among forms. Inverse dynamics solutions such as the one utilized in this study do not explicitly calculate moment arms. AnyBody has implemented a method of back-calculating moment arms and we provide this information for all lower limb muscle elements and the three degrees of rotation about the hip for both the modern human and australopithecine-like ADL models in spreadsheet formatted files. Moment arms are calculated for each muscle element moving through an idealized range of motion: 20 deg of extension to 90 deg of flexion; 30 deg of adduction to 50 deg of abduction; 40 deg of internal rotation to 40 deg of internal rotation.

#### Additional muscle activation plots

As with Figure 5 in the main text, all trials for the ADL human-like configuration are shown with blue dashed lines. The average for the ADL human-like (blue) and ADL australopithecine-like (gold) are shown with solid lines.

**Group A** muscles arise on the pelvis and insert on the femur proximal to the lesser trochanter (or proximal to the femoral shaft). They include (AnyBody muscle identifier): Iliacus, ObturatorExternus, ObturatorInternus, QuadratusFemoris, Piriformis, GemellusInferior, GemellusSuperior, GluteusMaximus, GluteusMedius, GluteusMinimus. The gluteal muscle activations are shown in Figure 5. Plots for the other Group A muscles are shown below.

**Group B** muscles cross the hip joint but arise either superiorly to the pelvis (PsoasMajor) or insert distally to the lesser trochanter. This latter set includes: AdductorBrevis, AdductorLongus, AdductorMagnus, BicepsFemorisCaputLongum, Sartorius, RectusFemoris, Gracilis, Semitendinosus, Semimembranosus, TensorFasciaeLatae. Plots for the Group B muscles are shown below.

**Group C** muscles are lower limb muscles, but do not cross the hip. They include: BicepsFemorisCaputBreve, ExtensorDigitorumLongus, ExtensorHallucisLongus, FlexorDigitorumLongus, FlexorHallucisLongus, GastrocnemiusLateralis, GastrocnemiusMedialis, Pectineus, PeroneusBrevis, PeroneusLongus, Plantaris, Popliteus, SoleusLateralis, SoleusMedialis, TibialisAnterior, TibialisPosterior, VastusIntermedius, VastusLateralis, VastusMedialis. We do not provide plots of muscle activation for these muscles.

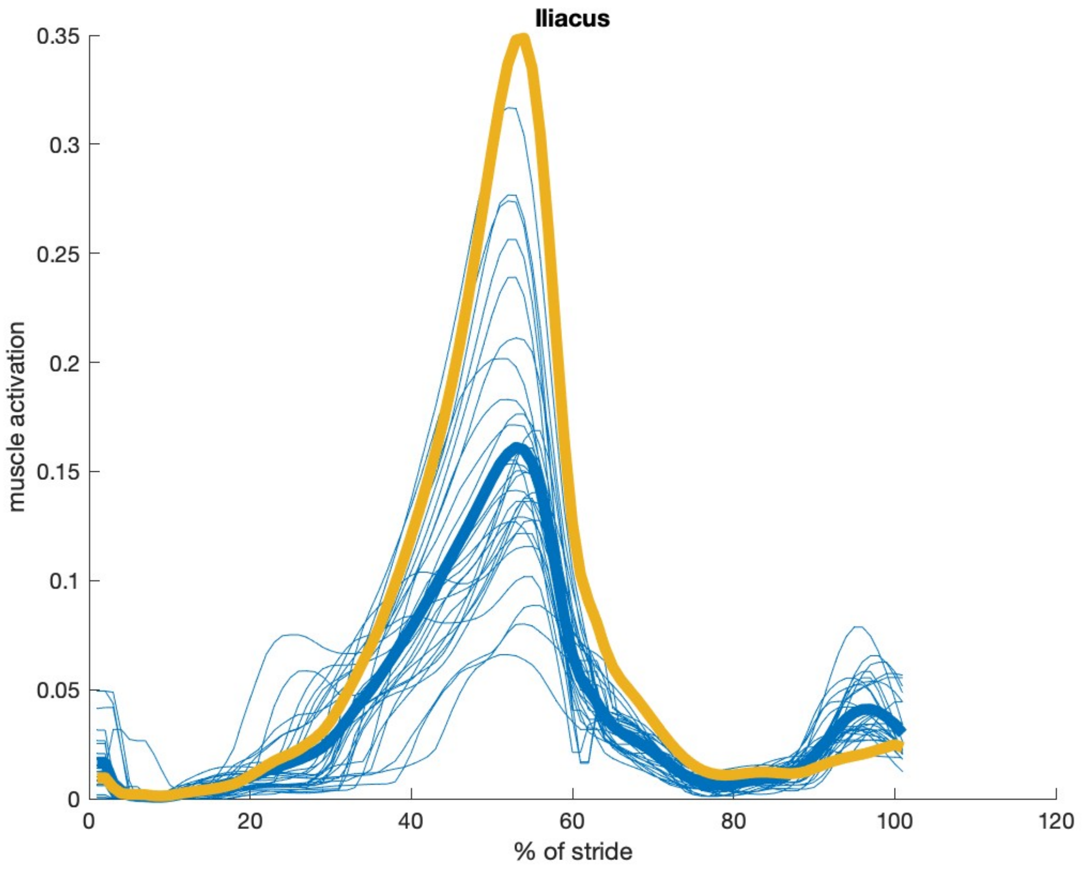
Iliacus
muscle activation
% of stride
0
0.05
0.1
0.15
0.2
0.25
0.3
0.35
0
20
40
60
80
100
120

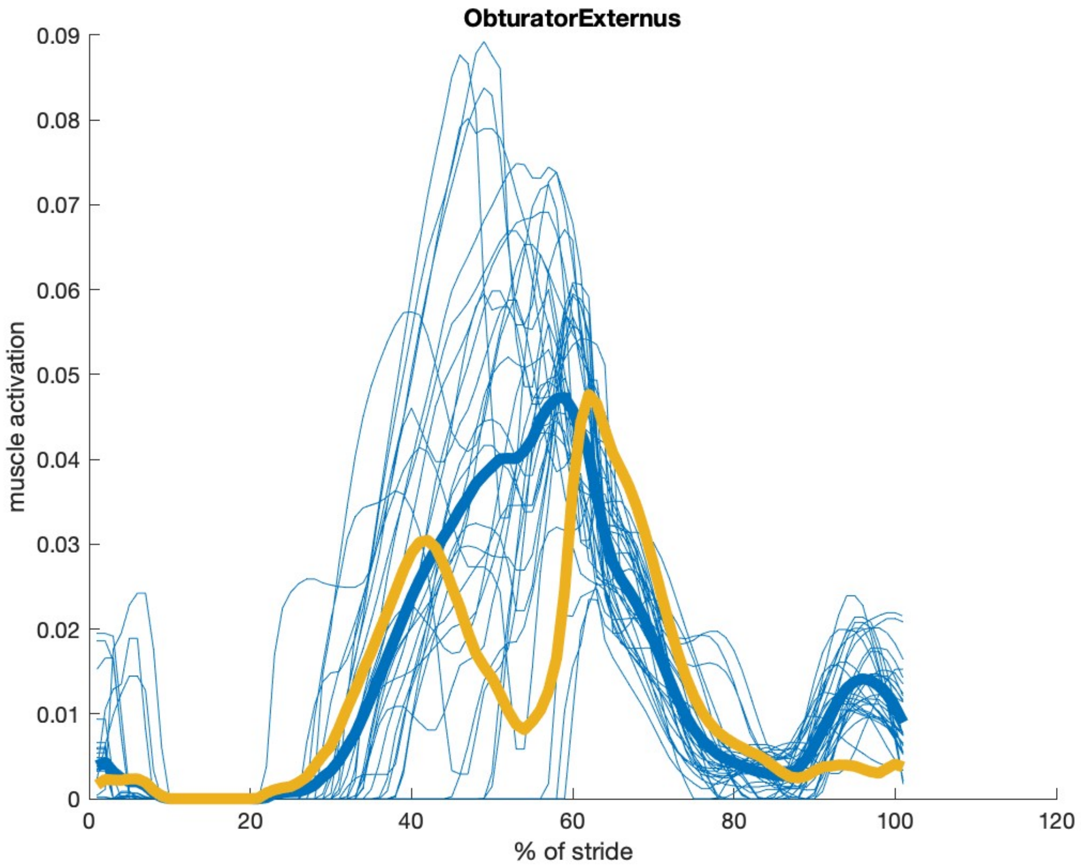

ObturatorExternus
muscle activation
% of stride
0.09
0.08
0.07
0.06
0.05
0.04
0.03
0.02
0.01
0
20
40
60
80
100
120

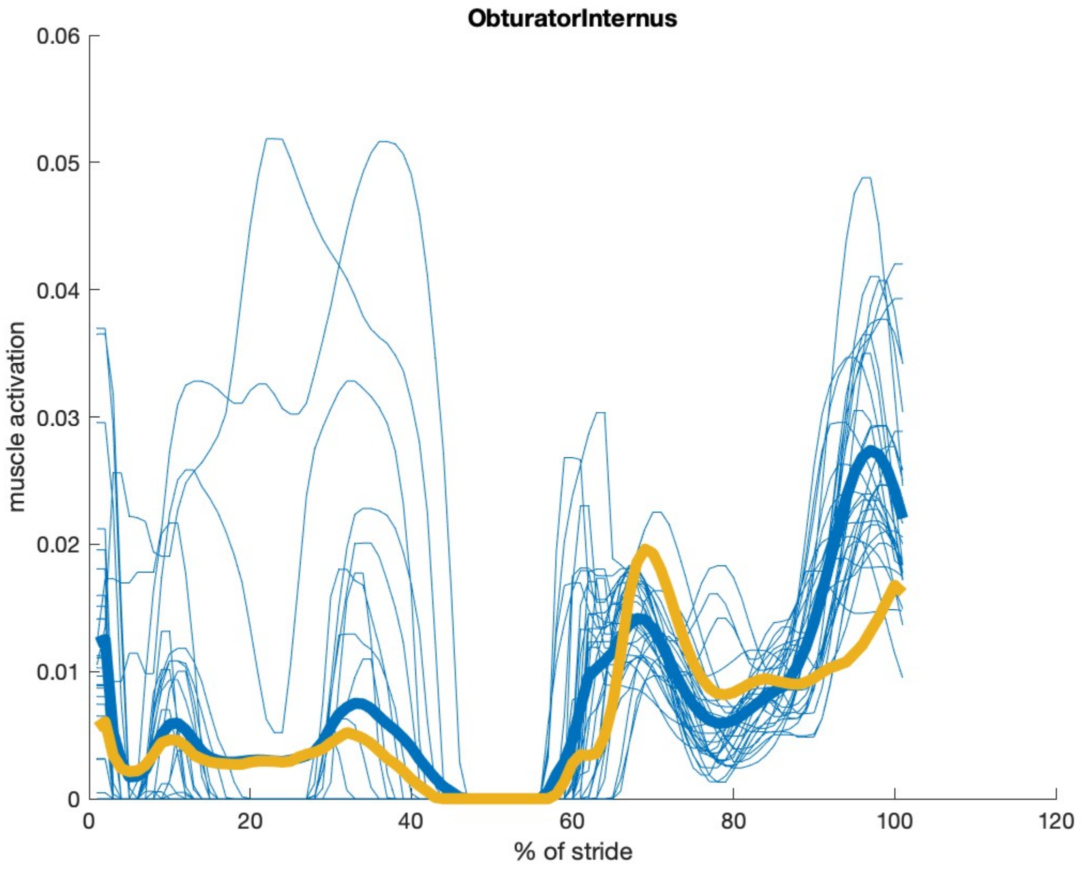
ObturatorInternus
muscle activation
% of stride
0.06
0.05
0.04
0.03
0.02
0.01
0
0
20
40
60
80
100
120

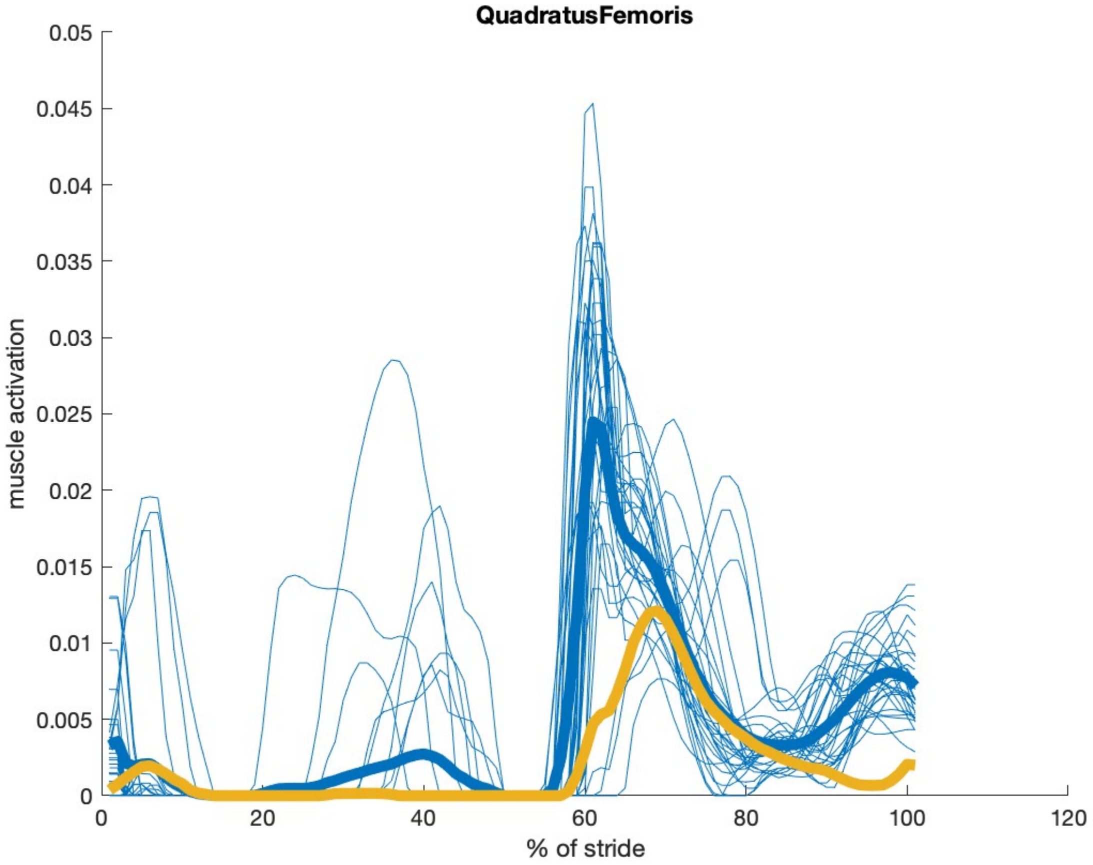

QuadratusFemoris
muscle activation
% of stride
0.05
0.045
0.04
0.035
0.03
0.025
0.02
0.015
0.01
0.005
0
0
20
40
60
80
100
120

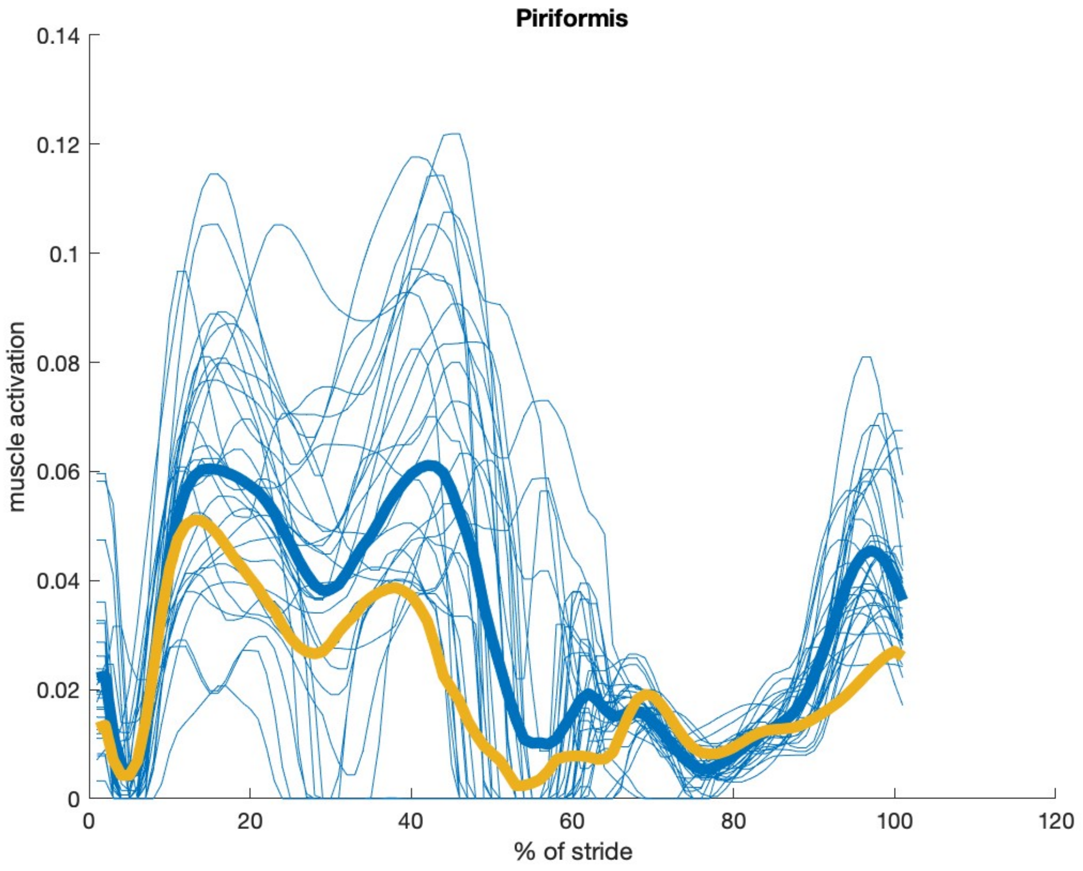

Piriformis
0.14
0.12
0.1
0.08
0.06
0.04
0.02
0
muscle activation
0
20
40
60
80
100
120
% of stride

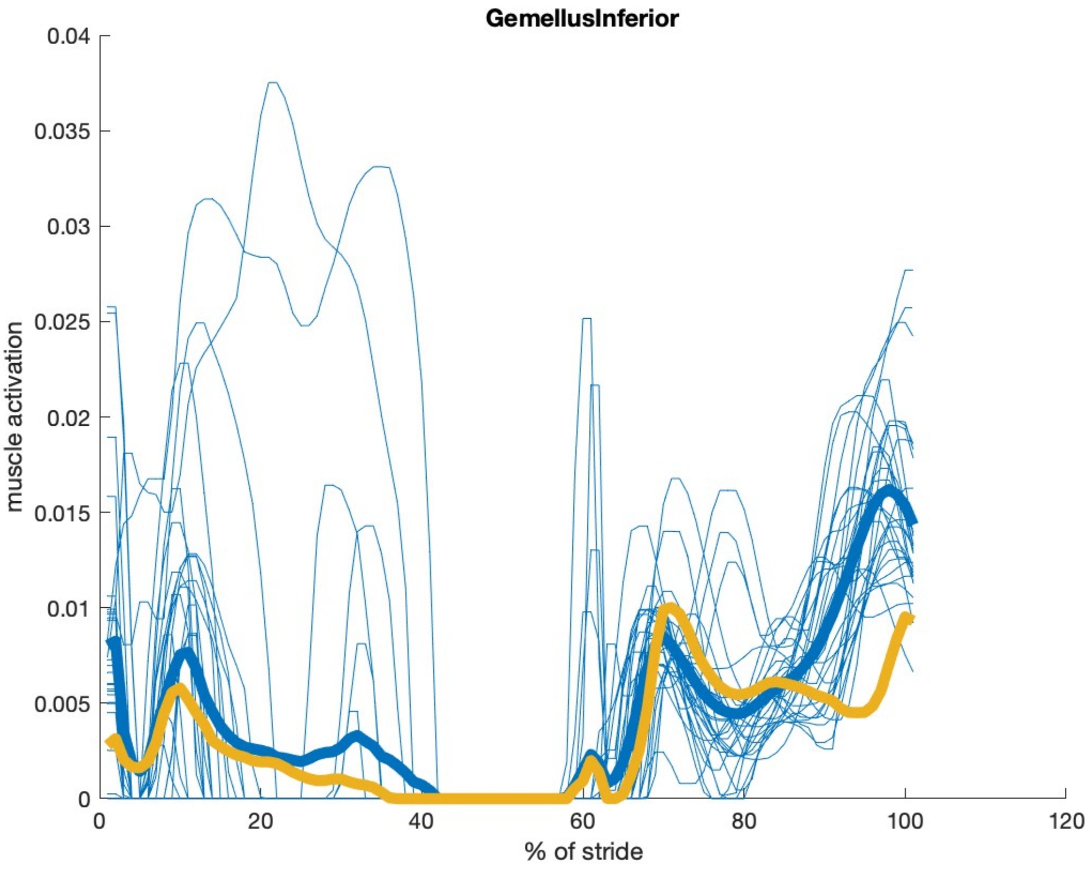
GemellusInferior
muscle activation
% of stride
0.04
0.035
0.03
0.025
0.02
0.015
0.01
0.005
0
0
20
40
60
80
100
120

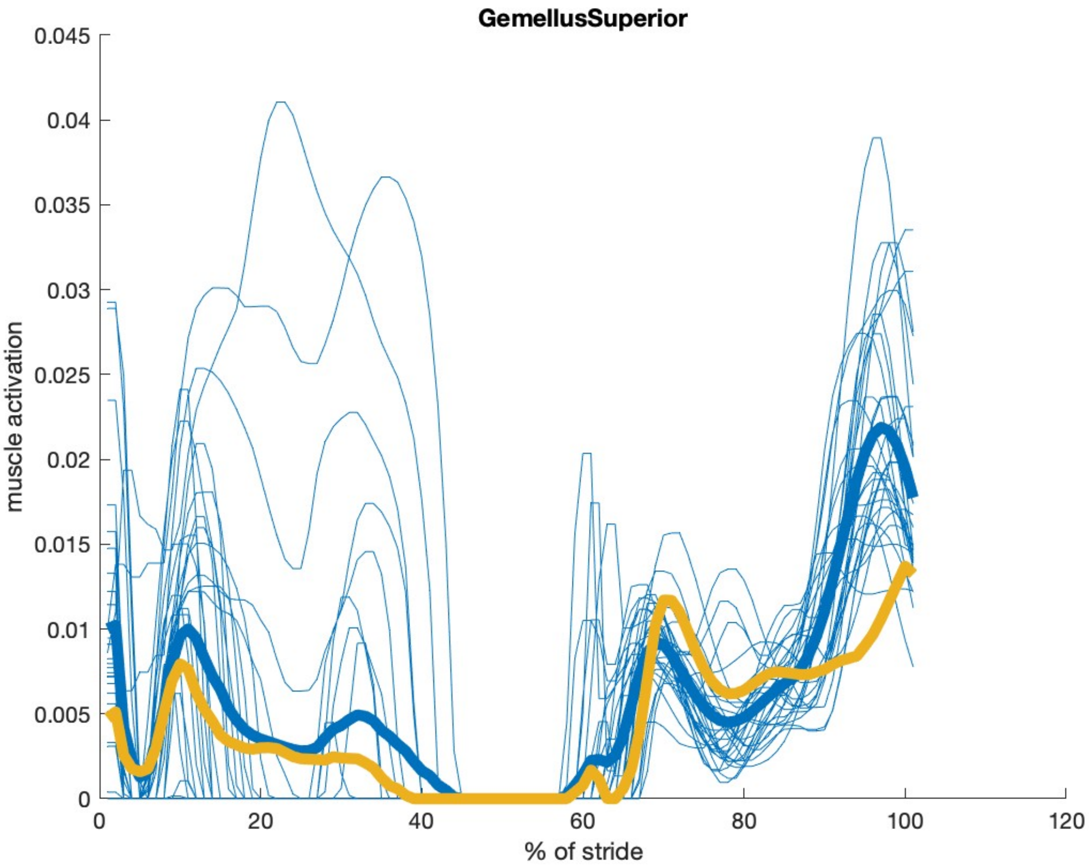

GemellusSuperior
muscle activation
% of stride
0.045
0.04
0.035
0.03
0.025
0.02
0.015
0.01
0.005
0
0
20
40
60
80
100
120

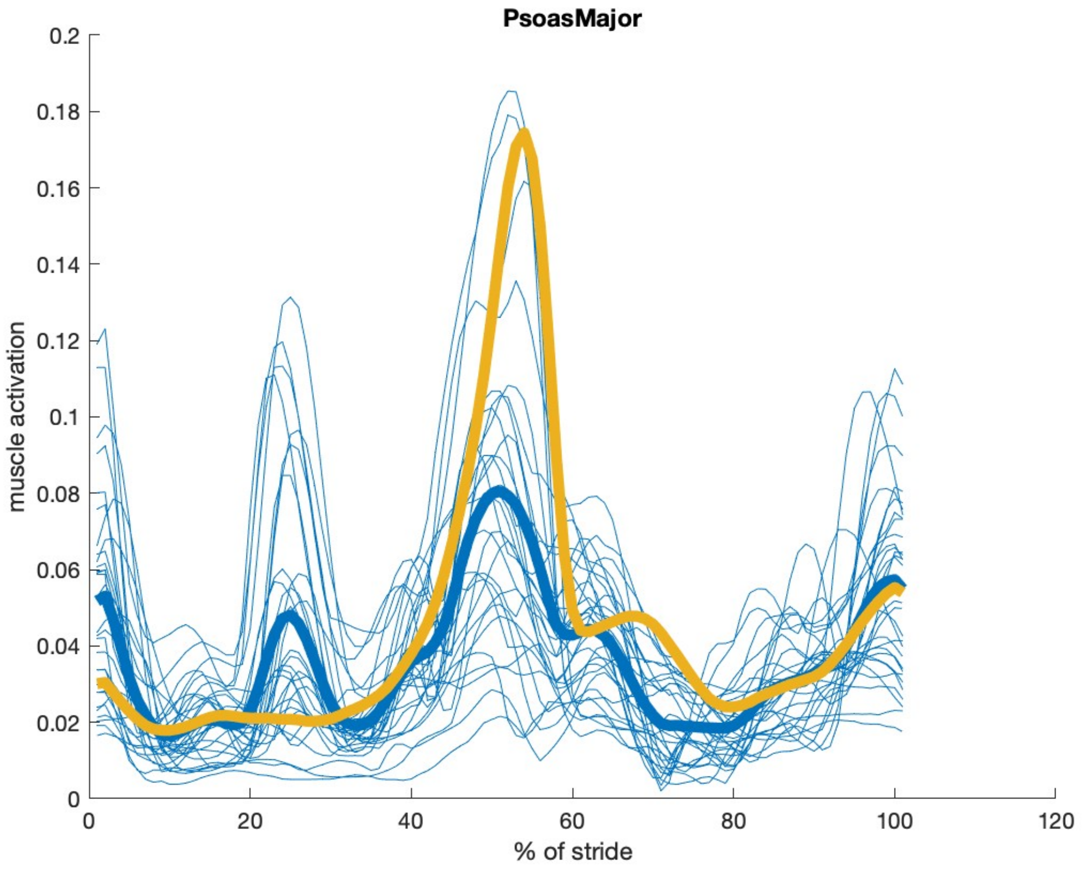

PsoasMajor
muscle activation
% of stride
0.2
0.18
0.16
0.14
0.12
0.1
0.08
0.06
0.04
0.02
0
0
20
40
60
80
100
120

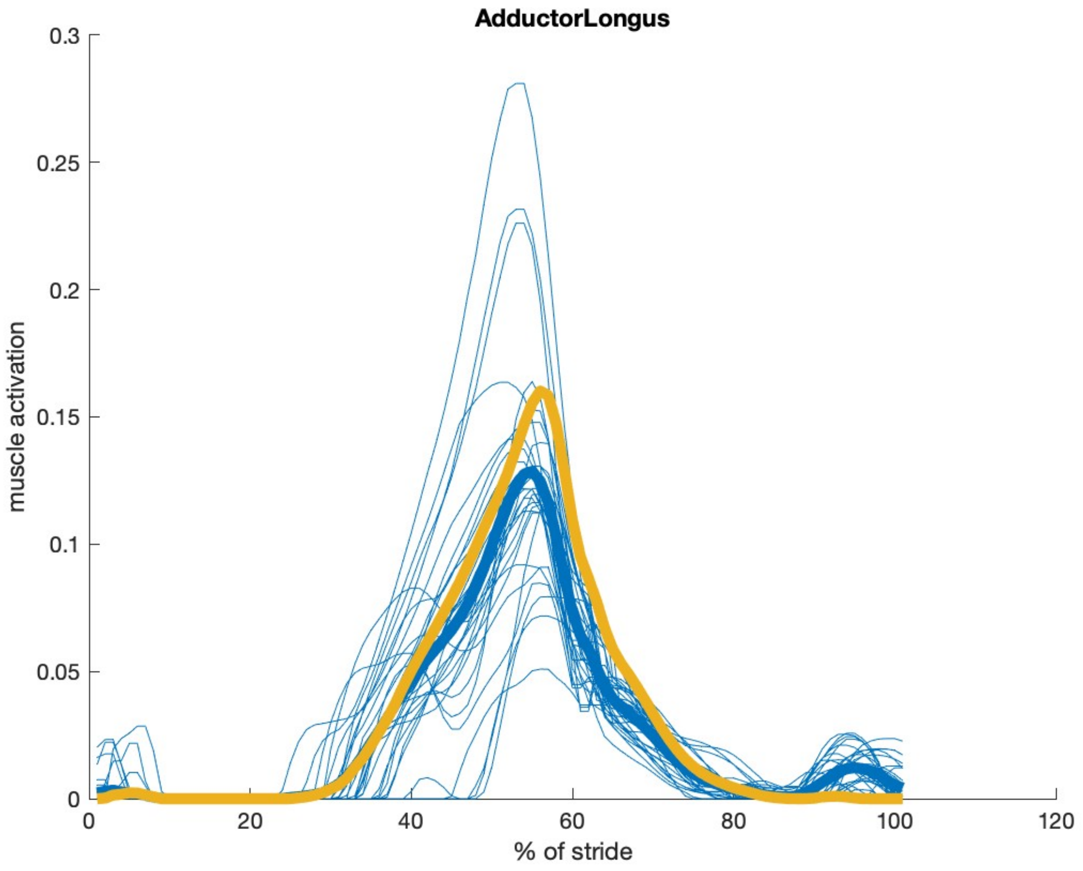
AdductorLongus
muscle activation
% of stride
0
0.05
0.1
0.15
0.2
0.25
0.3
0
20
40
60
80
100
120

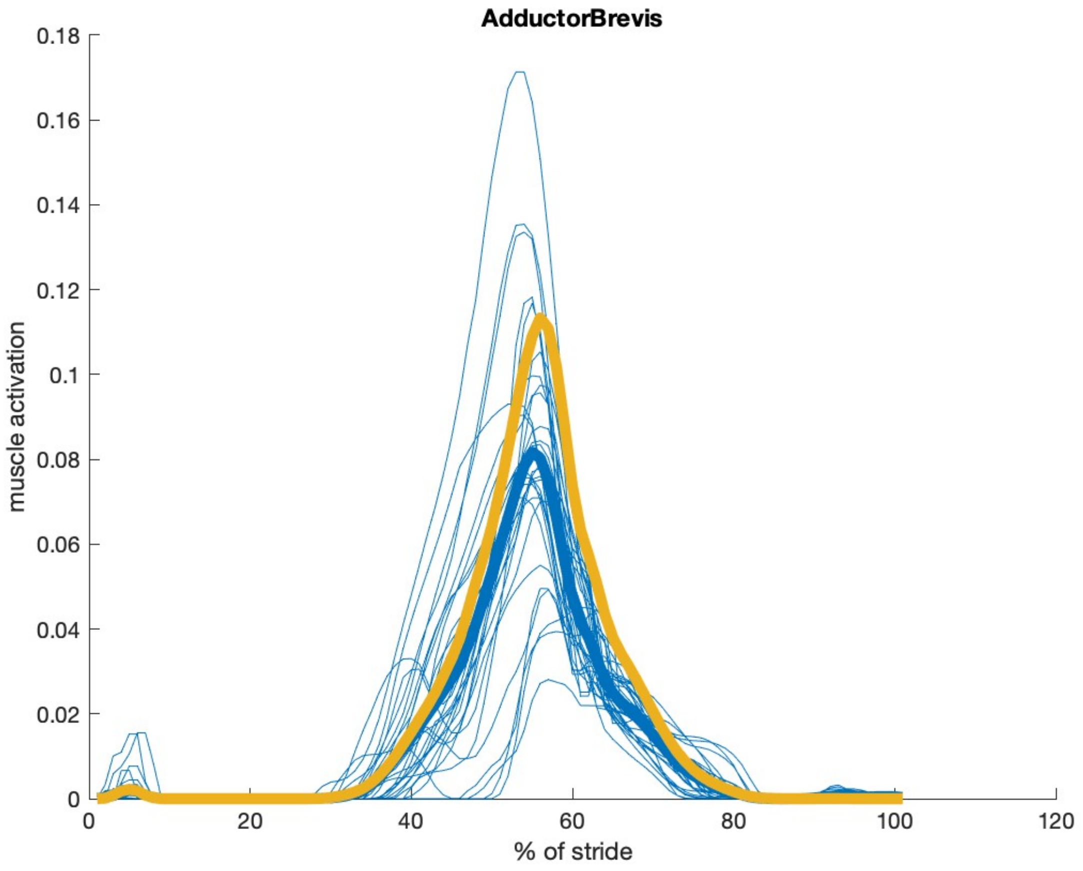

AdductorBrevis
muscle activation
% of stride
0.18
0.16
0.14
0.12
0.1
0.08
0.06
0.04
0.02
0
0
20
40
60
80
100
120

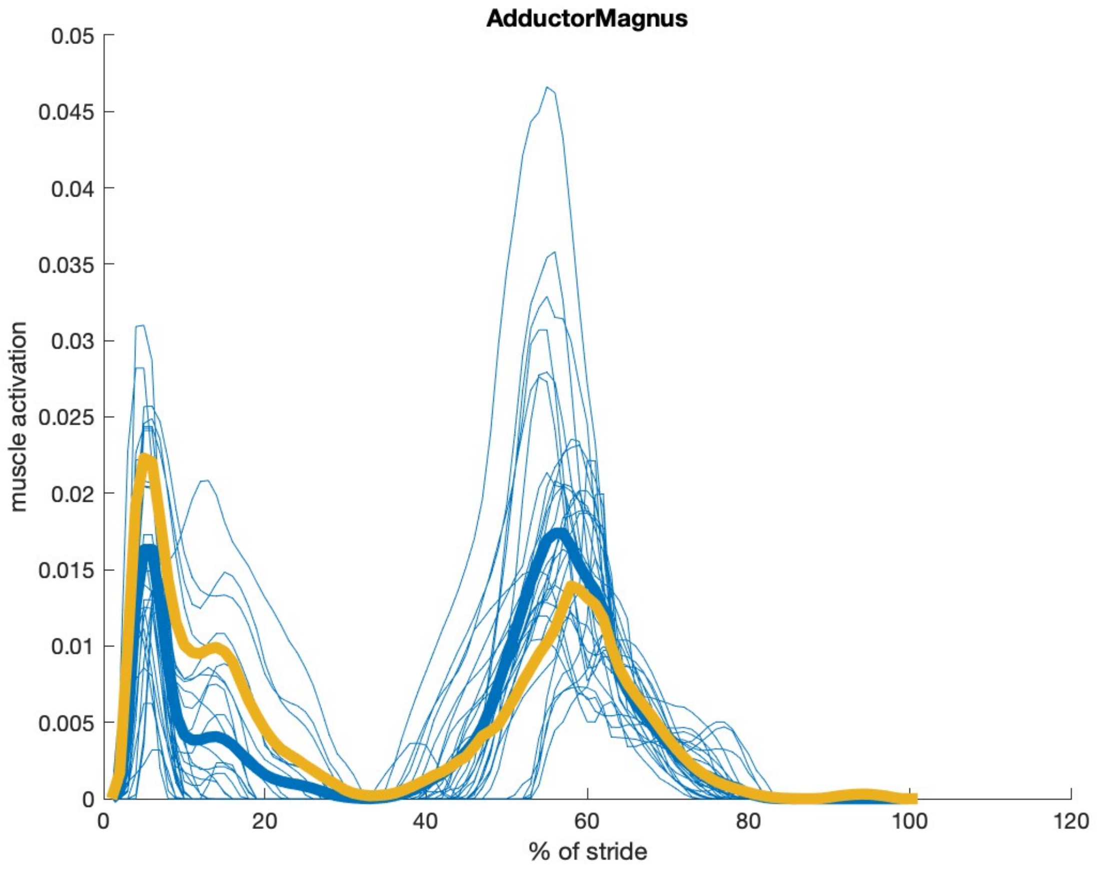

AdductorMagnus
muscle activation
0.05
0.045
0.04
0.035
0.03
0.025
0.02
0.015
0.01
0.005
0
0
20
40
60
80
100
120
% of stride

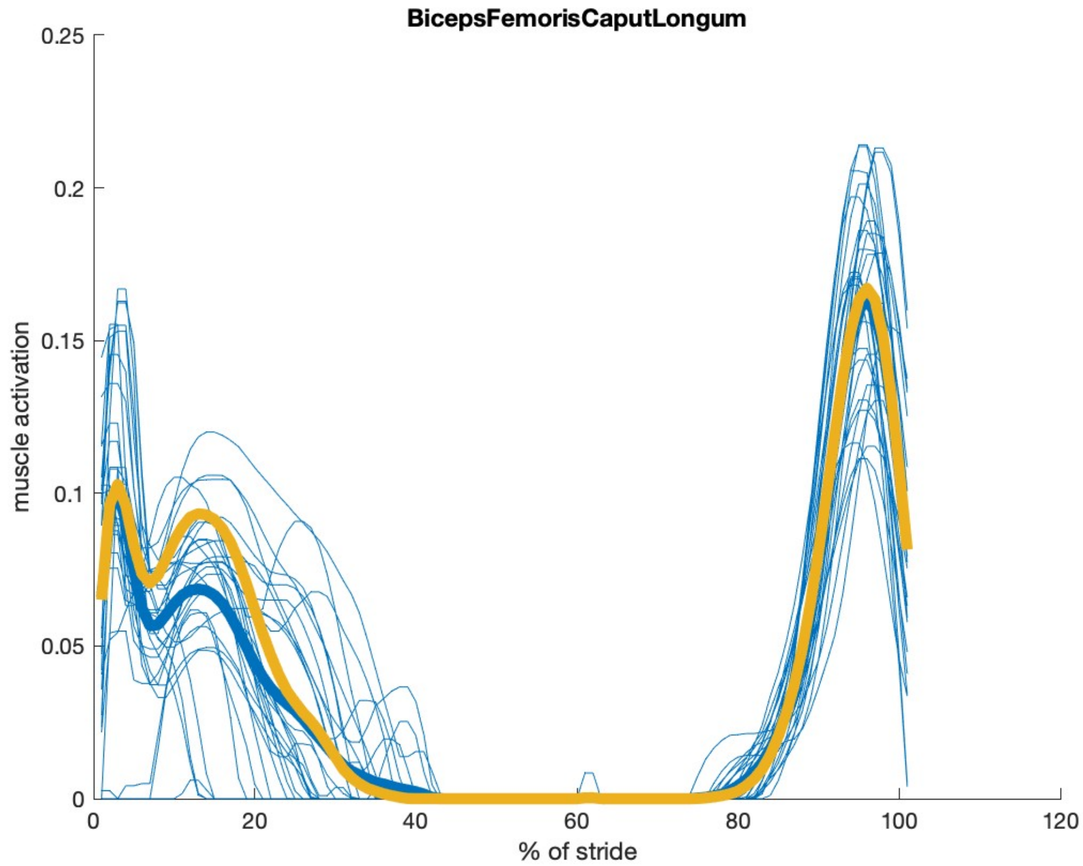

BicepsFemorisCaputLongum
muscle activation
% of stride
0
0.05
0.1
0.15
0.2
0.25
0
20
40
60
80
100
120

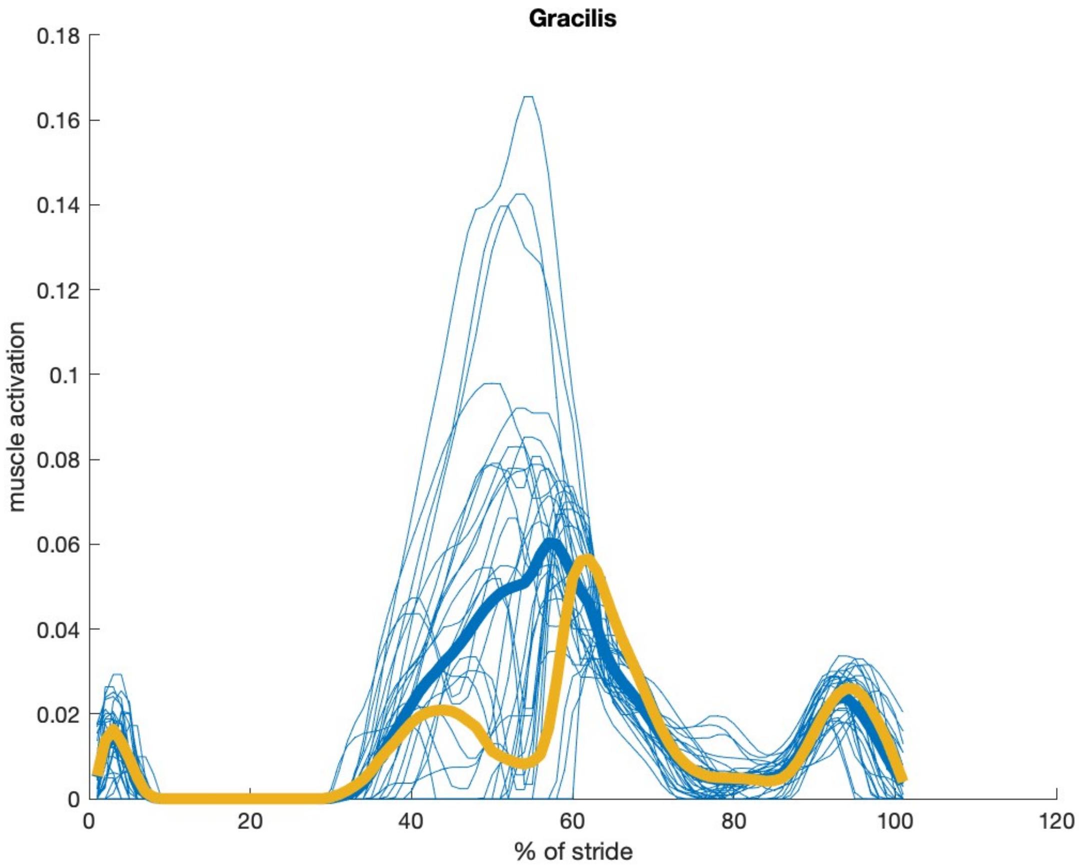

Gracilis
muscle activation
% of stride
0.18
0.16
0.14
0.12
0.1
0.08
0.06
0.04
0.02
0
20
40
60
80
100
120

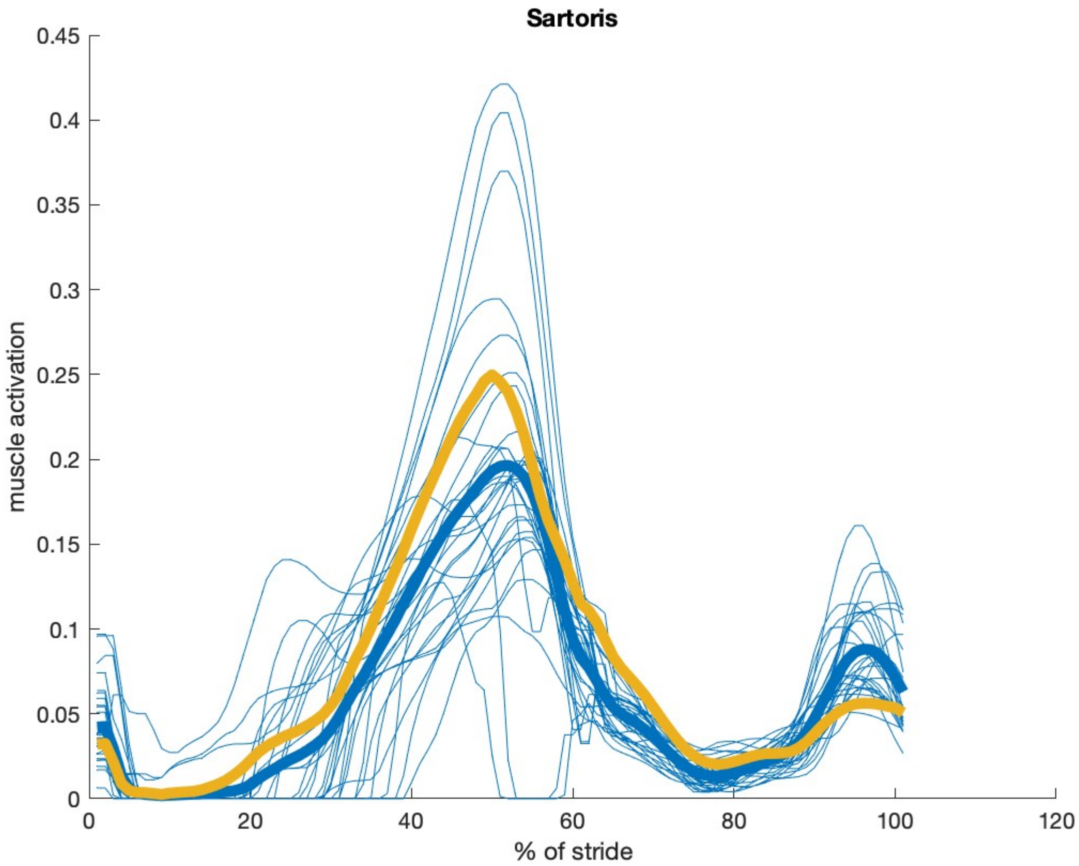

Sartoris
0.45
0.4
0.35
0.3
0.25
0.2
0.15
0.1
0.05
0
muscle activation
0
20
40
60
80
100
120
% of stride

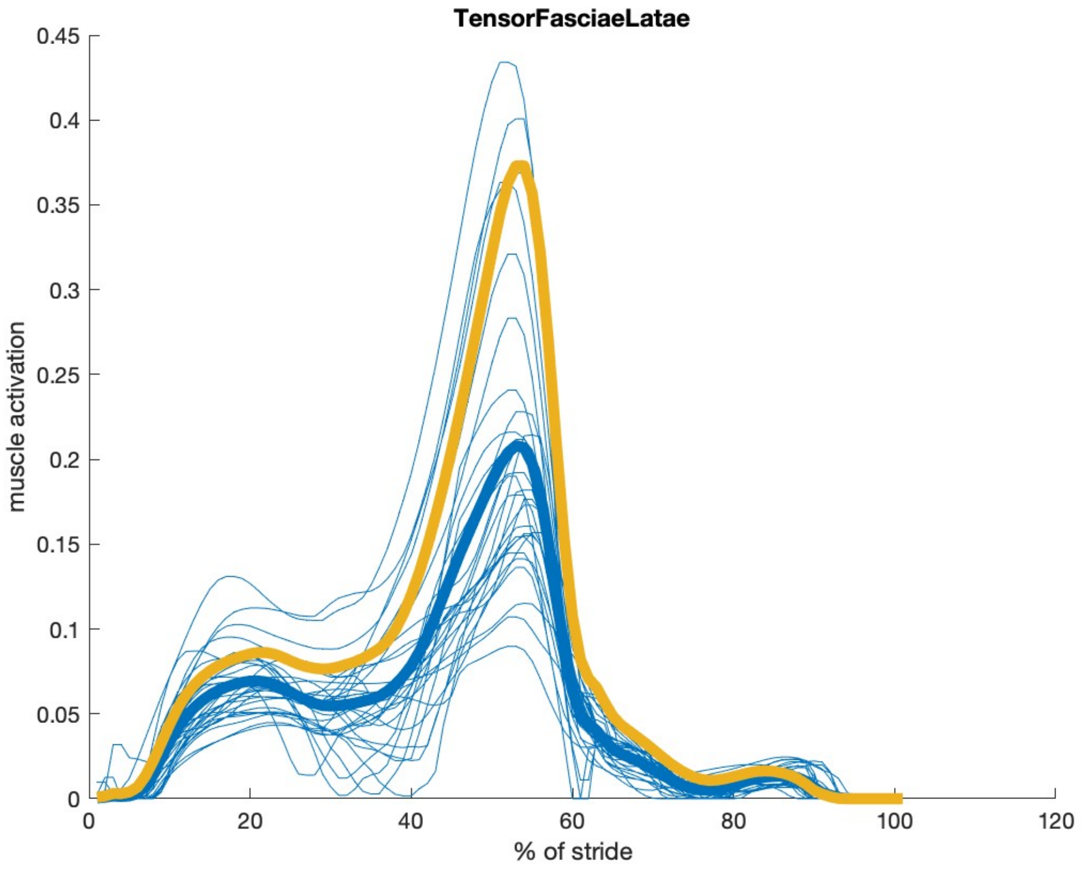

TensorFasciaeLatae
muscle activation
0.45
0.4
0.35
0.3
0.25
0.2
0.15
0.1
0.05
0
0
20
40
60
80
100
120
% of stride

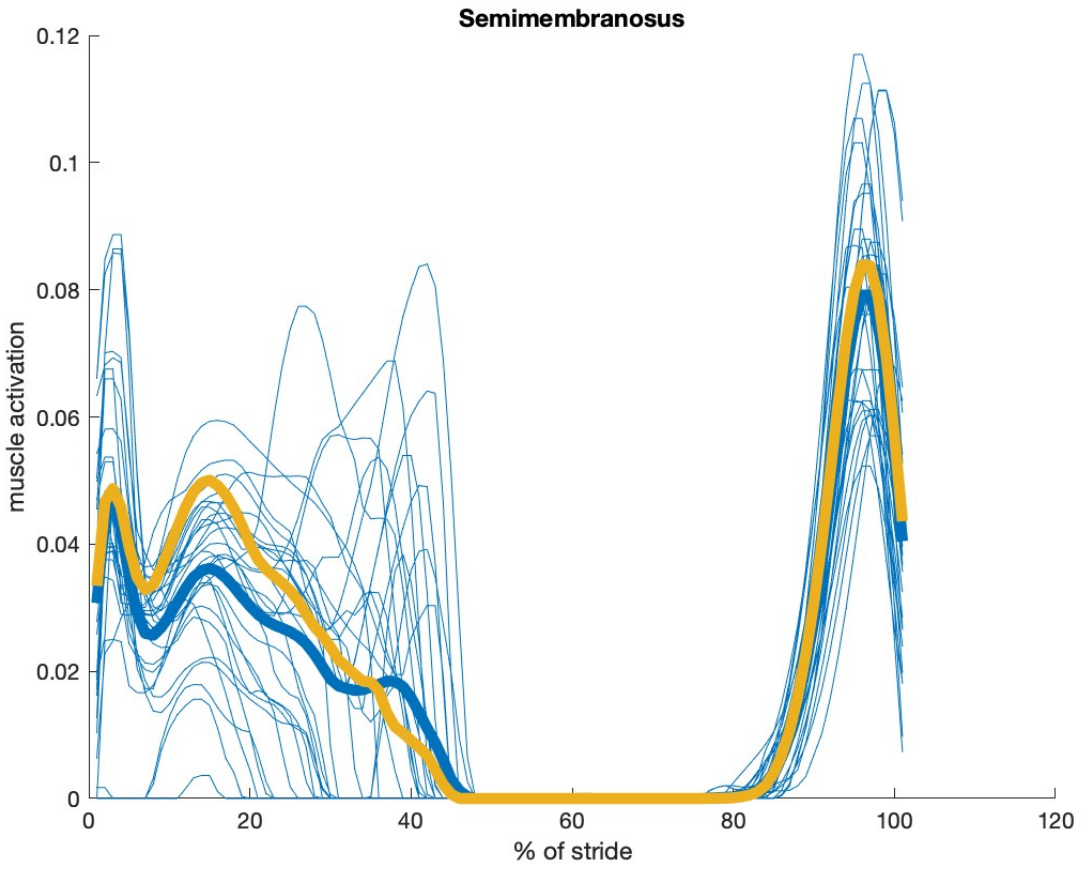
Semimembranosus
0.12
0.1
0.08
0.06
0.04
0.02
0
muscle activation
0
20
40
60
80
100
120
% of stride

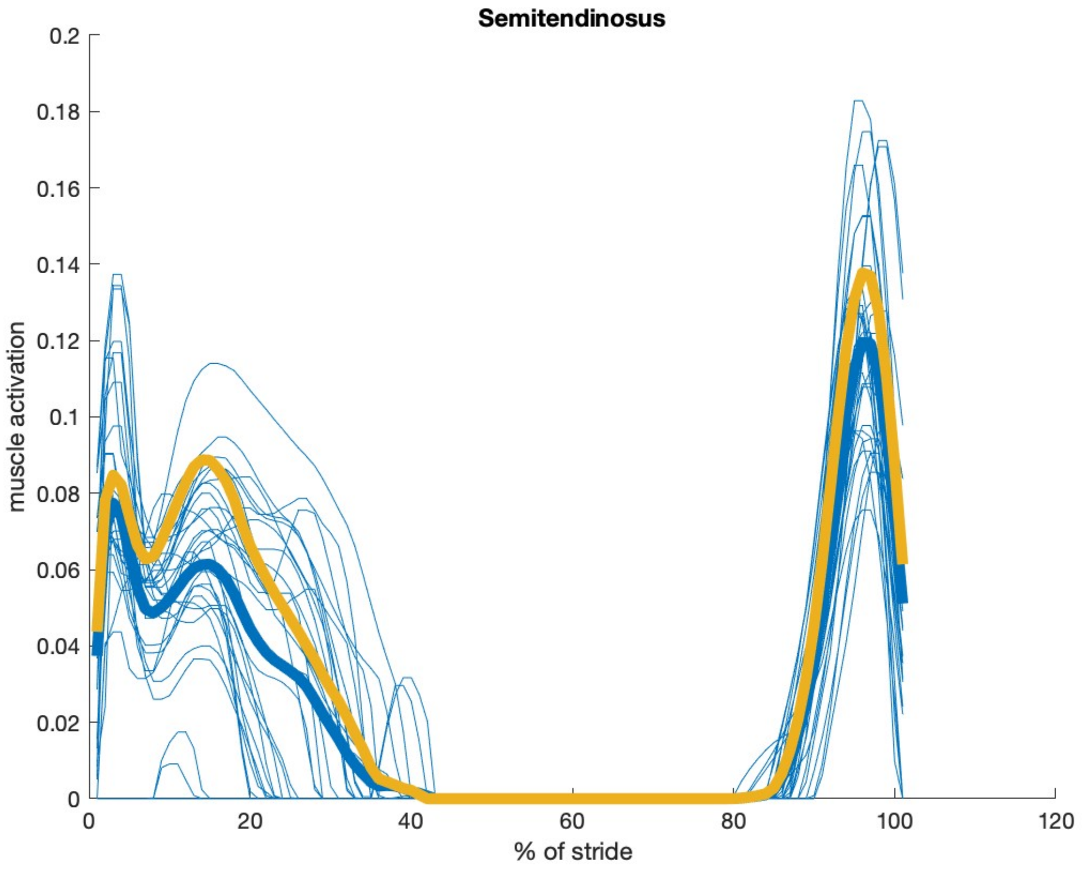
Semitendinosus
muscle activation
% of stride
0.2
0.18
0.16
0.14
0.12
0.1
0.08
0.06
0.04
0.02
0
0
20
40
60
80
100
120

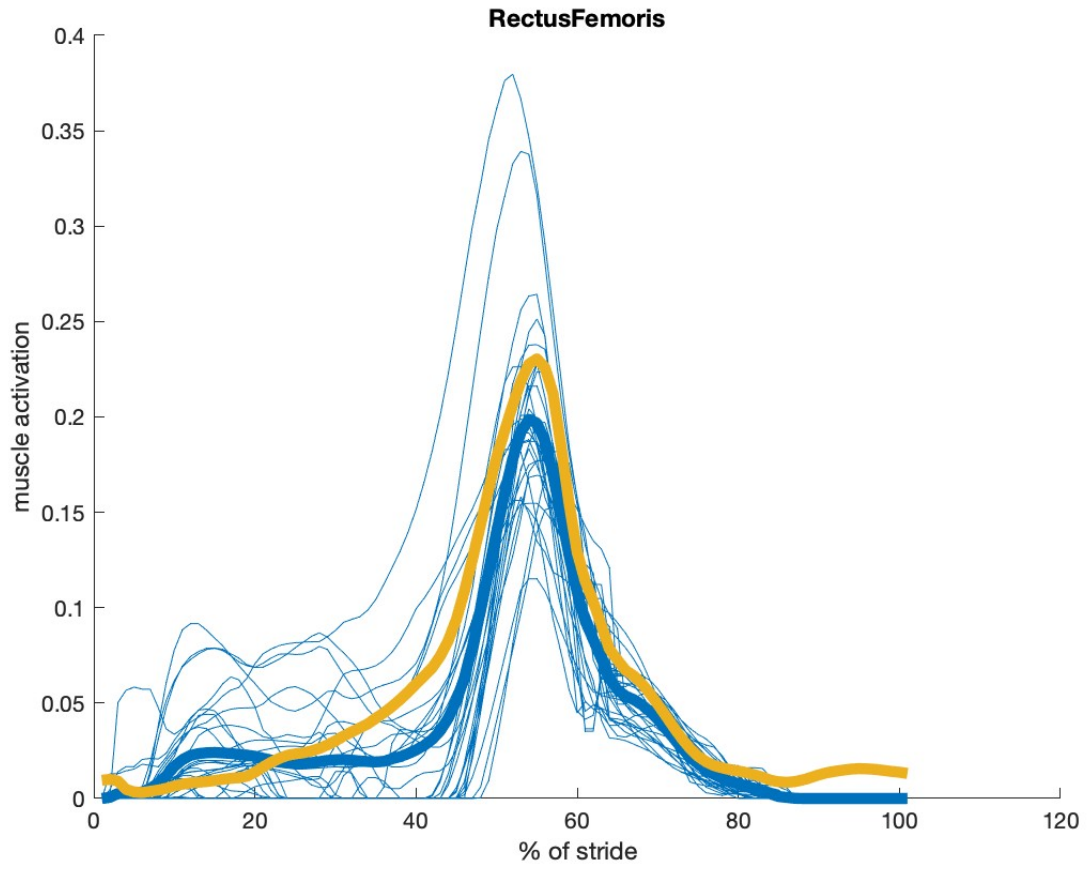
RectusFemoris
muscle activation
% of stride
0
0.05
0.1
0.15
0.2
0.25
0.3
0.35
0.4
0
20
40
60
80
100
120
